# Fibroblastic reticular cells provide a supportive niche for lymph node-resident macrophages

**DOI:** 10.1101/2021.11.24.469855

**Authors:** Joshua D’Rozario, Konstantin Knoblich, Mechthild Lütge, Christian Pérez Shibayama, Hung-Wei Cheng, Yannick O. Alexandre, David Roberts, Joana Campos, Jillian Astarita, Emma Dutton, Muath Suliman, Alice E. Denton, Shannon J. Turley, Richard L. Boyd, Scott Mueller, Burkhard Ludewig, Tracy Heng, Anne L Fletcher

## Abstract

The lymph node (LN) is home to resident macrophage populations that are essential for immune function and homeostasis. The T cell paracortical zone is a major site of macrophage efferocytosis of apoptotic cells, but key factors controlling this niche are undefined. Here we show that fibroblastic reticular cells (FRCs) are an essential component of the LN macrophage niche. Macrophages co-localised with FRCs in human LNs, and murine single-cell RNA-sequencing revealed that most reticular cells expressed master macrophage regulator CSF1. Functional assays showed that CSF1R signalling was sufficient to support macrophage development. In the presence of LPS, FRCs underwent a mechanistic switch and maintained support through CSF1R-independent mechanisms. These effects were conserved between mouse and human systems. Rapid loss of macrophages and monocytes from LNs was observed upon genetic ablation of FRCs. These data reveal a critically important role for FRCs in the creation of the parenchymal macrophage niche within LNs.

## Introduction

In lymph nodes, stromal cell communication with leukocytes is key to the initiation of a healthy immune response and eventual pathogen control (Fletcher *et al.*, 2020; Lütge, Pikor and Ludewig, 2021). Fibroblastic reticular cells (FRCs) are the most prevalent non-hematopoietic cell type in lymph nodes. Together with sinusoidal vascular elements, FRCs create the structural scaffolding on which leukocytes migrate and interact, including the T cell and dendritic cell-rich paracortex, the cortical B cell follicles, and the medullary plasma cell niche (Cremasco *et al.*, 2014; Huang *et al.*, 2018; Fletcher *et al.*, 2020). From this unique position at the coalface of the immune response, FRCs have evolved an immune-specialised role in regulating the survival, interaction, migration and function of T cells, B cells, dendritic cells, plasma cells, and innate lymphoid cells (ILCs) (Acton *et al.*, 2012; Cremasco *et al.*, 2014; Huang *et al.*, 2018; Knoblich *et al.*, 2018; Fletcher *et al.*, 2020; Pikor *et al.*, 2020; Kapoor *et al.*, 2021; Lütge, Pikor and Ludewig, 2021).

The importance of FRCs in adaptive immunity is well established, and emerging evidence suggests that they also play a critical role in innate immunity. Specialised Gremlin1^+^ subsets of fibroblasts engage in multifaceted crosstalk with dendritic cells in lymphoid tissues, and in some systems fibroblasts and macrophages are capable of co-operative support through mutual provision of growth factors (Zhou *et al.*, 2018; Bellomo *et al.*, 2021; Franklin, 2021; Kapoor *et al.*, 2021). Mouse FRCs also respond to viral and bacterial ligands (Fletcher *et al.*, 2010; Malhotra *et al.*, 2012; Yang *et al.*, 2014), regulate activation of group 1 ILCs and viral clearance through expression of the innate immunological sensing adaptor MyD88 (Gil-Cruz *et al.*, 2016), and in nonclassical secondary lymphoid organs, FRC-dependent MyD88 signaling steers B cell responses via TNF-dependent interactions with inflammatory monocytes (Perez-Shibayama *et al.*, 2018).

Despite the pivotal role of macrophages as managers of immune homeostasis and drivers of humoral and anti-viral immunity, our understanding of macrophage biology in lymph nodes is still evolving. Macrophages are found in every FRC niche (Baratin *et al.*, 2017; Bellomo *et al.*, 2018), but key factors controlling their development, function and localisation in lymph nodes remain unclear. Lymph node macrophages are categorized by the niches they occupy. Sinus-lining macrophages are relatively well-studied as highly phagocytic “flypaper” macrophages bathed in lymph and capable of engulfing incoming antigen for efficient presentation to other cell types (Bellomo *et al.*, 2018). Subcapsular sinus macrophages capture particles or immune complexes for direct transfer to follicular B cells, and are important for limiting viral spread (Carrasco and Batista, 2007; Junt *et al.*, 2007; Phan *et al.*, 2009; Gonzalez *et al.*, 2010; Iannacone *et al.*, 2010). They can present antigen to naïve T cells, but their ability to internalise and process antigen is lower than medullary sinus macrophages, which are highly efficient at pathogen and apoptotic cell clearance (Phan *et al.*, 2009). Depletion of both subsets in mice reduced anti-tumor immunity through a reduction in CD8^+^ T cell activation (Asano *et al.*, 2011). Mesenchymal lymphoid tissue organizer (LTO) cells and marginal reticular cells (MRC) utilize RANKL to support the development of subcapsular and medullary CD169^+^ sinusoidal macrophages, but not other macrophage subsets (Camara *et al.*, 2019).

Less well studied are the macrophages found in parenchymal and FRC-rich regions of lymph nodes. Medullary cord macrophages regulate medullary plasma cell maturation and survival (Mohr *et al.*, 2009; Huang *et al.*, 2018), while in the T cell paracortical zone, a rich, previously misidentified network of immunosuppressive macrophages play a unique role in immune homeostasis through potent efferocytosis of apoptotic T cells (Baratin *et al.*, 2017; Bellomo *et al.*, 2018). Both macrophage populations strongly co-localise with their local FRC network (Bellomo *et al.*, 2018; Huang *et al.*, 2018).

Here, we identified an essential support system provided by FRCs to macrophages. We demonstrate that expression of CSF1R ligands by FRCs is capable of regulating macrophage differentiation and survival. We further show in two genetic models of FRC ablation that depletion of FRCs drives a rapid loss of myeloid cells in lymph nodes. These data show that FRCs are a critically important component of the lymph node macrophage niche.

## Results

### T zone macrophages colocalize with FRCs in secondary lymphoid organs

Examination of mouse lymph nodes confirmed that macrophages identified by expression of MERTK co-localized with the FRC network in the T cell zone (**Figure 1A**) (Baratin *et al.*, 2017). We quantified these interactions by calculating macrophage proximity and alignment with extracellular matrix fibers secreted by FRCs, based on co-localization of >10% of macrophage perimeter with a visible fiber, for cells with a clear cross-section (**Figure 1B**). Macrophages were significantly more likely to be associated with an FRC fiber than not associated (**Figure 1C**), and FRC-associated macrophages showed significantly greater elongation (**Figure 1D**), supporting a physical association between these two cell types.

**Figure 1.**
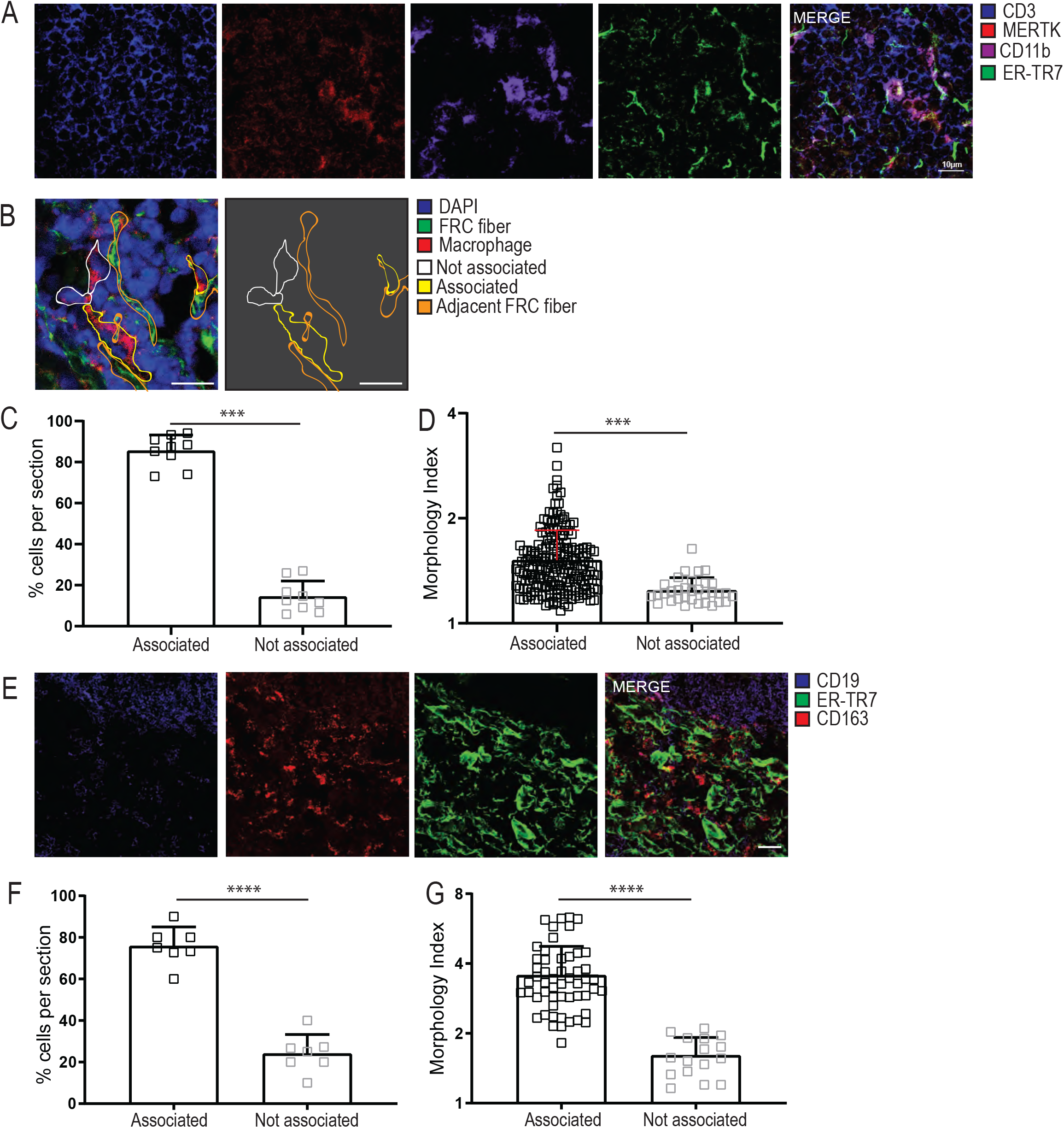
T zone macrophages colocalize with FRCs. **A.** Mouse lymph node (LN) immunofluorescence depicting MERTK^+^CD11b^+^ macrophages and ERTR7^+^ FRCs within the T-zone region of the lymph node (CD3^+^), representative of n=3 mice. Scale bar = 10μm. **B.** Macrophages with a clear DAPI^+^ nucleus had their perimeter outlined, and were designated as either associated (yellow outline) or not associated (white outline) with ERTR7+ FRC reticular fibers based on the proportion of colocalised perimeter. Human tonsil section shown, representative of analysis for both mouse and human tissues. Scale bar = 25μm. **C.** The proportion of macrophages per mouse LN section designated as associated or not associated with ERTR7^+^ reticular fibers. Each datapoint represents an individual tissue section. Data depict n=3 mice. **D.** Morphology index (perimeter^2^ / (4π*area)) was calculated for each MERTK+ macrophage observable in a clear cross-section with visible DAPI^+^ nucleus. Each datapoint represents an individual macrophage, and aggregate data is shown from 224 cells, from n=3 mice and 8 tissue sections. Scale bar = 10μm. **E.** Human tonsil immunofluorescence depicting association of CD163^+^ macrophages with ER-TR7^+^ FRCs in an area of paracortex adjacent to a B cell follicle. Scale bar = 50μm. Imaging represents n=3 human donors. **F.** T-zone macrophages were designated as associated or not associated with ERTR7^+^ reticular fibers, as described in B. **G.** Morphology index was calculated for each CD169^+^ macrophage as described in C. Data is shown from 69 cells, from N=3 human donors and 7 tissue sections. All statistics shown are Mann-Whitney U test; *** P<0.001, **** P<0.0001.

Next, we examined whether the same held true for human FRCs and macrophages in tonsils. CD163^+^ macrophages co-localized with the FRC network in the paracortical T cell zone (**Figure 1E)**, and also preferentially associated with FRC fibers, with significant elongation when associated (**Figure 1F, G**).

These data provide evidence for a close relationship between macrophages and the FRC network that is conserved across human and mouse lymph nodes.

### Conserved expression of genes crucial for myeloid cell maturation, migration and function

To determine whether FRCs express factors relevant to myeloid cell maintenance, and whether these were broadly concordant across species, we performed transcriptomic analysis and comparison of a range of FRC datasets obtained from cultured human tonsil-derived FRC samples (**Figure 2A**), cultured versus freshly isolated human tonsil-derived FRCs (**Figure 2B**), freshly isolated FRCs from mouse skin-draining (SLN) or mesenteric lymph nodes (MLN) (first published in 31) (**Figure 2C**) and cultured mouse SLN FRCs (**Figure 2D**). All datasets showed robust expression of macrophage colony stimulating factor CSF1 and myeloid chemoattractants CCL2, CXCL1, and CXCL12. CXCL8 has no mouse equivalent but was strongly expressed in human cells. Tonsil and SLN, but not MLN FRCs, showed high expression of monocyte differentiation and growth factor IL-6. These data showed that expression of myeloid regulatory factors is a property that is conserved across human and mouse FRCs, and suggested a potential role for the FRC-macrophage dyad in regulating the myeloid response to infection.

**Figure 2:**
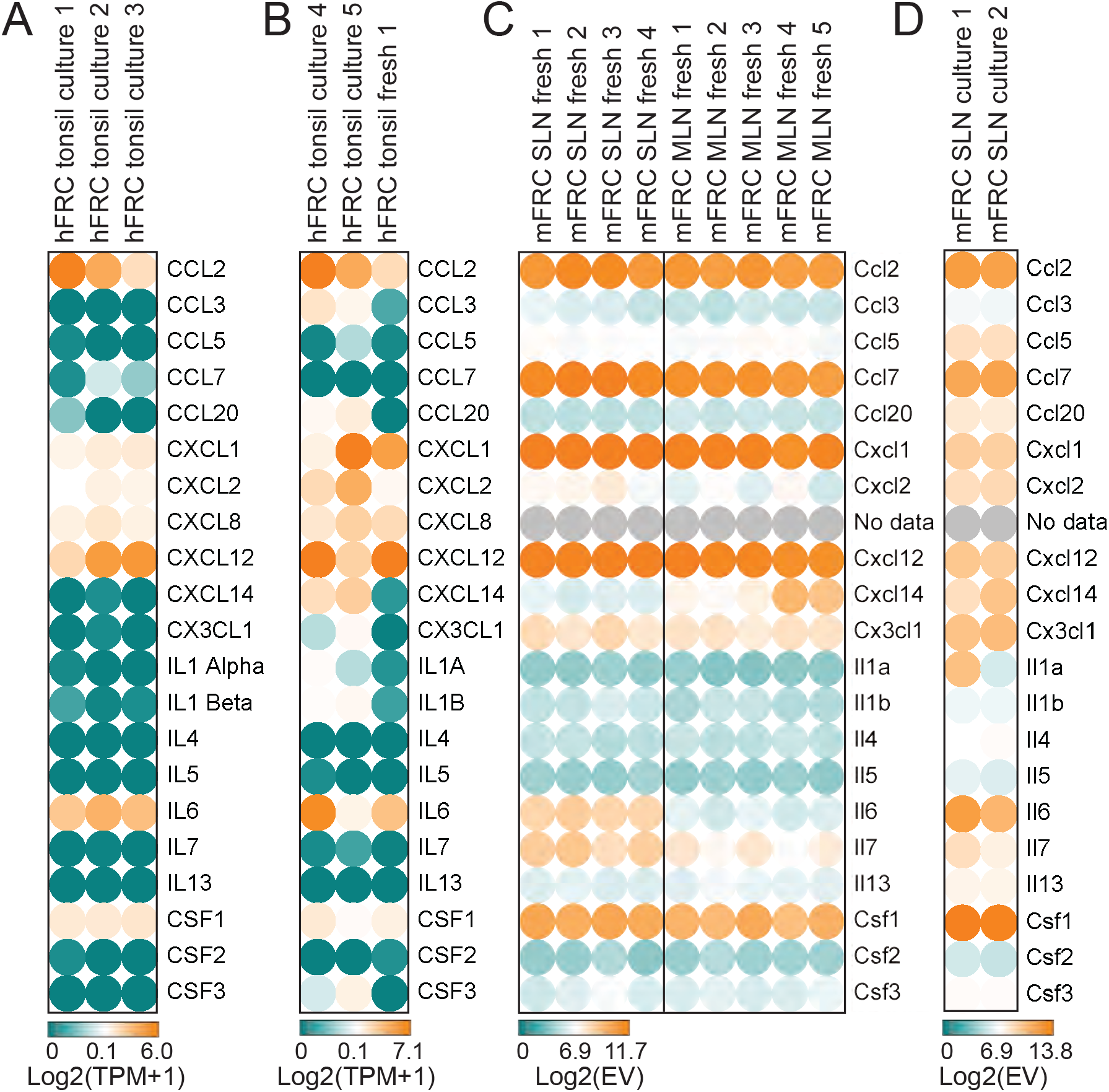
Mouse and human FRCs express genes relevant to myeloid cell maturation, recruitment and function. **A.** RNA-Seq data from culture-expanded human tonsil FRCs harvested at passage 3. n=3 unrelated human donors are depicted per group. **B.** RNA-Seq data from 2 cultured and 1 freshly isolated human tonsil FRCs. n=3 unrelated human donors are depicted per group Donors from B and A are unrelated. **C.** Microarray data from freshly isolated mouse lymph node FRCs, obtained from pooled donors. n=4 individual datasets for skin-draining lymph nodes and n=5 individual datasets for mesenteric lymph nodes. **D.** Microarray data from cultured mouse skin-draining lymph node FRCs; samples obtained from n=2 individual datasets. Heatmaps were generated using Morpheus software (Broad Institute). EV = expression value. hFRC = human fibroblastic reticular cells. mFRC = mouse fibroblastic reticular cells. TPM = transcripts per million. Below the expression threshold is shown as green, genes at the expression threshold are shown as white, transitioning to orange for expression, with maximum gene expression set as the orange maximum. Different platforms were used to acquire these datasets; they are not suitable for quantitative cross-comparisons across datasets.

### FRCs exhibit TLR4 signalling and expression of myeloid cell immunoregulatory factors

FRCs express pattern recognition receptors TLR3 and TLR4, and can respond to cognate bacterial and viral stimuli or analogues (Fletcher *et al.*, 2010; Malhotra *et al.*, 2012; Gil-Cruz *et al.*, 2016; Severino *et al.*, 2017). However, the kinetics, signalling mechanisms and effects on downstream transcription are not well defined, particularly for human FRCs. We therefore examined the effect of pathogen sensing on FRC-derived factors relevant to myeloid cell migration and function.

RNA sequencing (RNA-Seq) analysis of cultured human FRCs showed that TLR4 expression was the highest and least variably expressed TLR between the 3 donors (**Figure 3A**).

**Figure 3:**
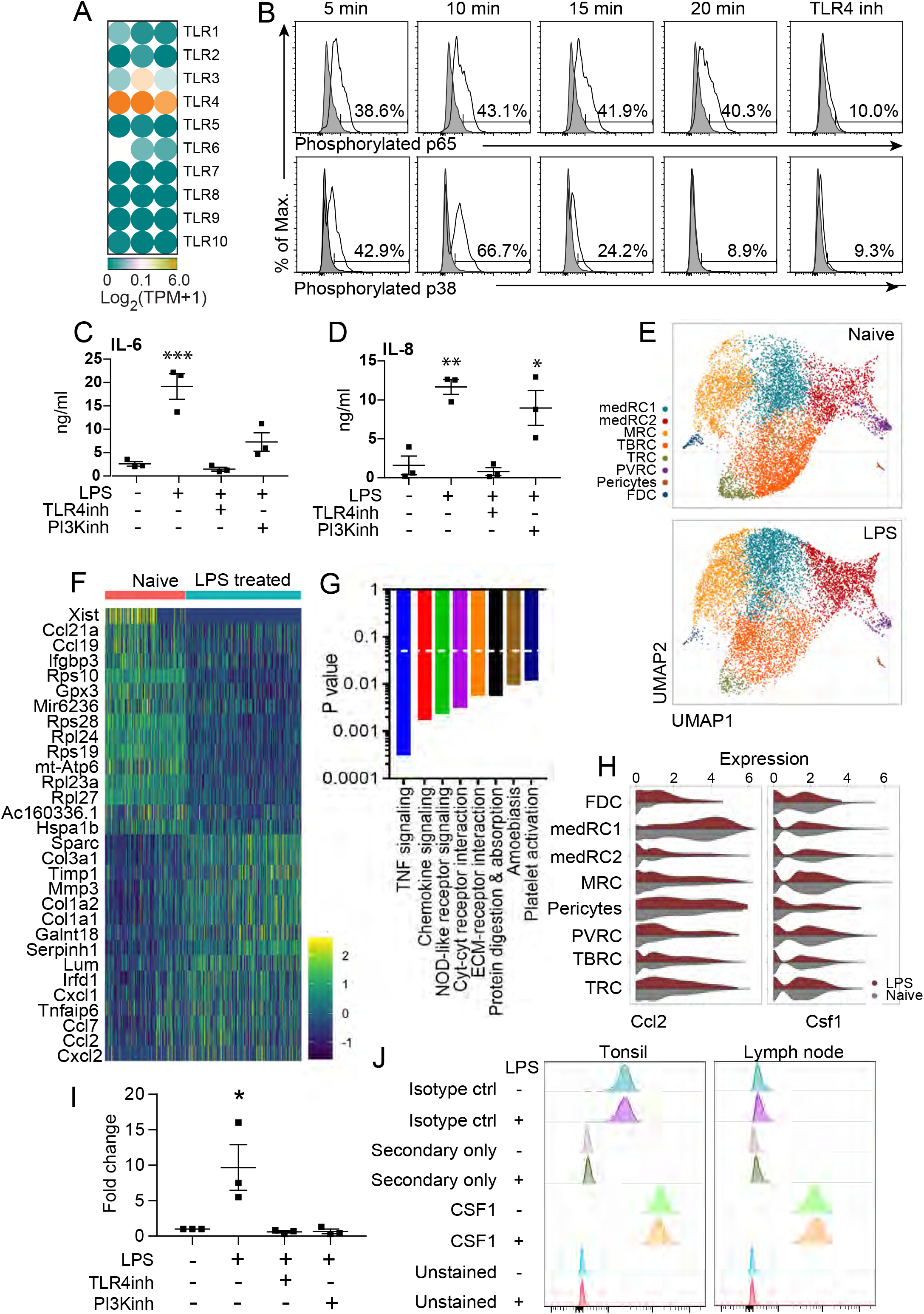
Human FRCs respond to TLR4 signaling. **A.** RNA-Seq heatmap data from cultured human FRCs showing TLR gene expression. TPM: transcripts per million. **B.** Human FRCs were cultured *in vitro* with or without TLR4 inhibitor CLI-095 as designated, then stimulated with 10 ng/ml of LPS, fixed over a 20 min time course, then analysed by flow cytometry for phosphorylation of p65 and p68. Unstimulated control shown in grey and used for multiple panels. Histograms represent 2-3 FRC donors from 6 independent experiments. **C, D:** Human lymph node or tonsil FRCs from 3 donors were cultured *in vitro* and stimulated with 1μg/ml of LPS for 24h, with cytokine quantities measured using Luminex Bead technology at ng/ml as shown. **E.** EYFP^+^ reticular cells were isolated from brachial lymph nodes from Ccl19 Cre R26R-EYFP mice, which were either treatment-naïve, or immunised with OVA/LPS. scRNA-seq was performed on EYFP^+^ cells. UMAP of EYFP+ lymph node reticular cells colored by the assigned subsets, with or without treatment. **F.** Top 15 differentially expressed genes between reticular cell clusters from naïve and LPS treated mice. **G**. KEGG pathway analysis of genes upregulated with LPS treatment (*P*<0.05 depicted with dotted line, FDR and Benjamini-Hochberg <0.05). **H.** Violin plots showing expression of Ccl2 and Csf1 in treated or naïve mice. **I.** Human lymph node or tonsil FRCs from 3 donors were cultured *in vitro* and stimulated with 1μg/ml of LPS for 24h, with CCL2 protein measured using Luminex Bead technology. Fold-change from untreated cells is depicted. Mean + SEM shown, n = 3 individual human donors from 2 independent experiments. **J.** Human lymph node or tonsil FRCs from 2 donors were cultured with or without LPS as described in C. Flow cytometry for CSF1 is shown compared with staining controls. Data depict 2 independent experiments. For C, D and I, Mean + SEM shown, n = 3 individual human donors from 2 independent experiments. * p<.05, ** p<0.01, *** p< 0.001, one-way ANOVA with Tukey’s multiple comparison test, comparing to untreated. PI3Kinh = PI3K inhibitor; TLR4inh = TLR4 inhibitor.

To confirm functional MyD88-dependent TLR4 signalling within human FRCs, we assessed phosphorylation of p65 (RelA, NF-kB pathway) and p38 (MAPK pathway) in response to LPS (**Figure 3B**) (Akira and Takeda, 2004). TLR4 inhibitor CLI-095 (TAK-242), which specifically binds the intracellular domain of the TLR4 receptor to block signalling (Kawamoto *et al.*, 2008), was used to confirm TLR4 dependence. Flow cytometric analysis showed rapid phosphorylation of both p38 and p65 in FRCs after 5 minutes of LPS exposure, peaking at 10 minutes, confirming that FRCs quickly sense LPS and respond via functional TLR4 signalling. Accordingly, LPS stimulation increased the secretion of downstream targets IL-6 and IL-8 (CXCL8) by cultured human FRCs within 24h (**Figure 3C, D)**.

Next, we investigated global transcriptional outcomes of TLR signalling to FRCs *in vivo*. CCL19-Cre x R26R-EYFP mice were injected with LPS to induce an inflammatory response. After 3 days, EYFP+ reticular cells from the lymph nodes of LPS treated and control mice were sorted for single-cell RNA-Seq analysis. Eight fibroblastic clusters were identified and validated via expression of expected markers (**Figure 3E; Supp. figure 1A-C**) (Rodda *et al.*, 2018; Pikor *et al.*, 2020). The subsets identified comprised T zone reticular cells (TRC); follicular dendritic cells (FDC); pericytes; perivascular reticular cells (PVRC); marginal reticular cells (MRC); T-B zone reticular cells (TBRC); and two subsets of medullary reticular cells (medRC1 and medRC2). (**Figure 3E).**) All clusters were represented in both treatment groups (**Figure 3E; Supp. figure 1D**).

Next, we compared LPS-treated versus treatment-naïve mice and identified the 30 most differentially expressed genes (15 upregulated with treatment, 15 downregulated) across all clusters. LPS treatment was associated with upregulated expression of myeloid cell-attracting chemokines including Cxcl1, Cxcl2 and Ccl2. Conversely, expression of homeostatic chemokines Ccl19 and Ccl21a was downregulated, as well as genes encoding ribosomal proteins. KEGG pathway analysis (*P*<0.05, FDR and Benjamini-Hochberg <0.05) showed that LPS treatment drove significant over-representation of genes related to innate immune function, including TNF, chemokine, cytokine and NOD-like receptor signalling (**Figure 3G**).

Based on their prominent roles in myeloid cell regulation, we chose to further validate expression of Ccl2 and Csf1, which were robustly expressed across all human and mouse RNA screening data (**Figure 2, 3**). Ccl2 was significantly upregulated with LPS treatment in mice, while Csf1 showed unchanged robust expression across fibroblast subsets (**Figure 3F, H**). To test if this held true in human cultures, FRCs from 3 donors were treated with LPS for 24h and secreted proteins analysed. LPS treatment drove a significant upregulation of CCL2 secreted protein, which was abrogated in the presence of TLR4 inhibitor TAK-242 (**Fig. 3I**). As the PI3K/AKT signaling pathway can be activated in MyD88-dependant TLR4 signalling (Laird *et al.*, 2009), we also examined the effect of PI3K inhibition of LPS-induced protein secretion by FRCs. In the presence of PI3K inhibitor LY294002, the increase in CCL2 was similarly abrogated (**Figure 3I**). These data confirmed that CCL2 upregulation by human FRCs was driven by LPS signaling through TLR4. Conversely, Csf1 expression did not appear as a differentially expressed gene in the mouse scRNA-Seq analysis, a finding borne out by flow cytometry staining for CSF1 protein expression in human FRCs, which was unchanged after LPS treatment (**Figure 3J**).

### Mouse and human FRCs can regulate the survival and differentiation of monocytes via signalling to CSF1R

A functional role for CSF1 expression by lymph node FRCs has not previously been reported. Robust expression of CSF1 across our mouse and human datasets, together with the close connection between FRCs and myeloid cells observed *in vivo*, led us to hypothesise that FRCs may have the capacity to support myeloid cell development or differentiation.

In the lymph nodes, T cell zone resident macrophages are long-lived, with slow replacement by blood-borne monocytes (Baratin *et al.*, 2017). Monocytes can be defined as classical (CD11b^+^Ly6C^hi^ in mouse, CD14^+^CD16^−^ in human) or non-classical (CD11b^+^Ly6C^lo^ in mouse, CD14^−/lo^CD16^+^ in human) based on their ability to perform a pro- or anti-inflammatory functions (Ziegler-Heitbrock *et al.*, 2010).

The proliferation, differentiation and survival of macrophages and monocytes is regulated by CSF1, which acts via autocrine or paracrine signaling through its receptor, CSF1R (Chitu and Stanley, 2006; Lenzo *et al.*, 2012). To investigate the effects of FRC-derived CSF1 signaling on myeloid cells, we performed co-culture assays with mouse bone marrow cells as a source of myeloid precursors. In the presence of recombinant CSF1, as expected, bone marrow cells differentiated to CD11b^+^F4/80^+^ macrophages (**Figure 4A**). This macrophage differentiation was inhibited by the addition of a CSF1R blocking antibody (**Figure 4A, Supp. Fig 2A**). Notably, the addition of FRCs to bone marrow cells, without exogenous CSF1, was sufficient to yield an expansion of (CD11B^+^ F4/80^+^) macrophages (**Figure 4A**).

**Figure 4:**
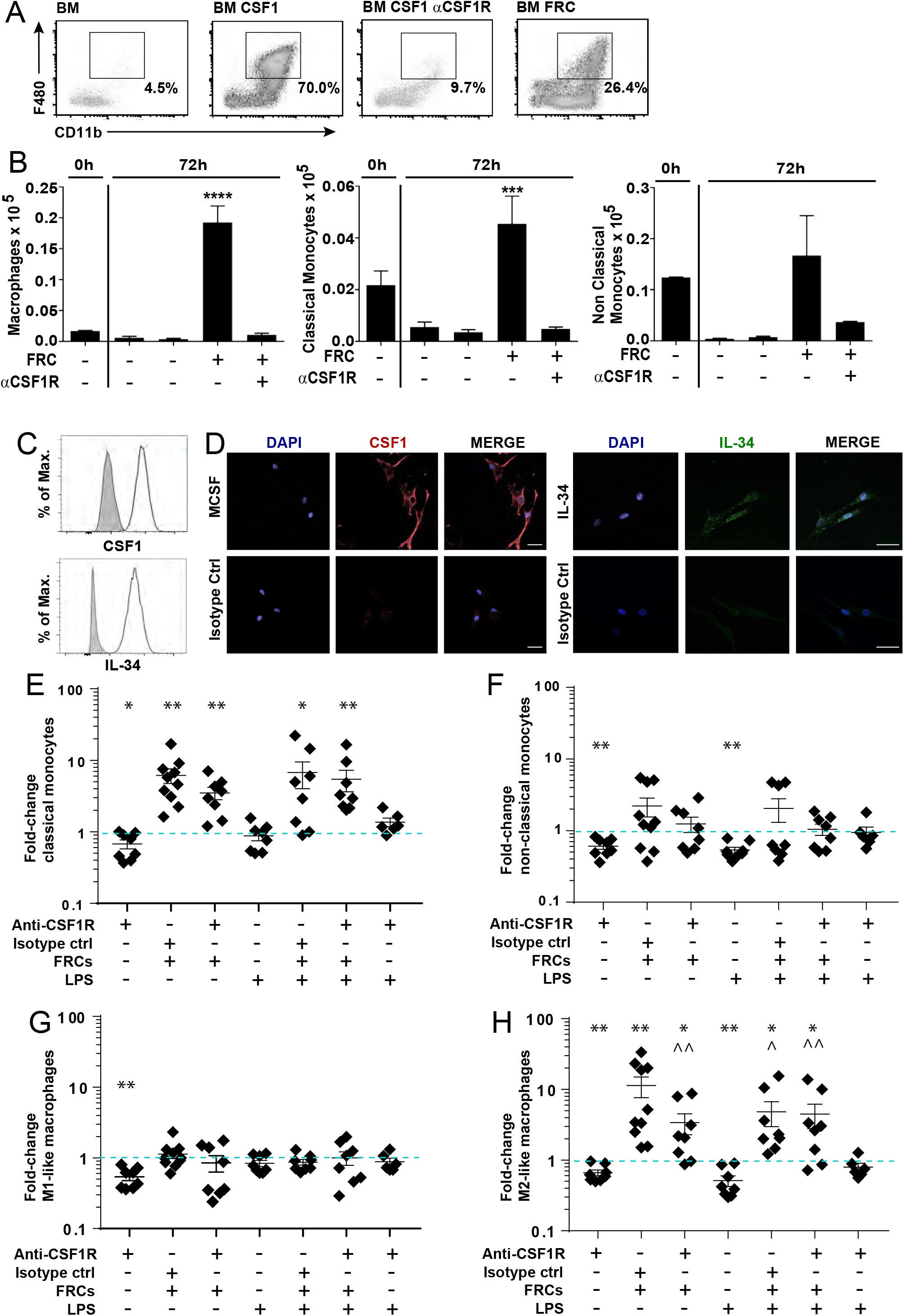
Fibroblastic reticular cells support monocyte differentiation via CSF1R signalling. **A, B**. 1×10^6^ mouse bone marrow cells were co cultured with 2×10^5^ mouse FRCs with or without LPS, TLR4 inhibitor (TLR4i), recombinant CSF1, or anti-CSF1R blocking antibody as shown. **A.** Representative flow cytometry profiles for F4/80 and CD11b after 72h of culture. Macrophages were gated on CD45^+^ GR-1^−^. Monocytes were gated as non-classical (GR-1^−^ CD11b^+^ Ly6C^lo^) or classical (GR-1^−^ CD11b^+^ Ly6C^hi^). **B.** Macrophage and monocyte numbers were quantified using flow cytometry at either 0h (input analysis) or after 72h of culture. Mean+SEM shown. Data depict n=4 mice, and are representative of 3 independent experiments. ***P<0.001, one-way ANOVA with Tukey’s post-test. Mean + SEM shown. **C.** Human FRCs at passage 3 were analysed for expression of IL-34 and CSF1 protein using flow cytometry. Isotype control antibody staining shown in gray, Representative of 2 donors. **D.** Immunofluorescent images of human FRCs stained for IL-34 and CSF1 protein compared to isotype control antibodies. Scale bars are 50μm. Representative of 2 donors. **E-H.** 4 × 10^5^ human peripheral blood mononuclear cells (PBMCs) were incubated with or without LPS, CSF1R blocking antibody, isotype control antibody, or 2×10^4^ human tonsil-derived FRCs. After 72 hours of culture, cells were quantified and analysed via flow cytometry. The fold change is compared to untreated controls (normalised to 1, shown as dotted blue line) is depicted for: **E.** M1-like macrophages (CD64^+^ HLA-DR^+^CD206^−^), **F:** M2-like macrophages (CD206^+^CD64^−^), **G:** Classical monocytes (CD16^−^CD14^+^), **H:** non-classical monocytes (CD16^+^CD14^−^). All subsets were gated negative for CD3, CD19, CD56, CD135. Data are representative of n = 2 FRC and n=3 PBMC donors from 3 experiments. Mean + SEM shown. * p<0.05, ** p<0.01; Wilcoxon rank test with a ratio paired T test. Star depicts significance vs untreated group (normalized to 1); chevron depicts significance vs FRC + isotype control group. FRCs: fibroblastic reticular cells. CSF1: colony stimulating factor 1; CSF1R: CSF1 receptor; LPS: lipopolysaccharide. αCSF1R: anti-CSF1R blocking antibody.

Next, bone marrow cells were co-cultured for 3 days with or without mouse FRCs or CSF1R blocking antibody (**Figure 4B**). The addition of FRCs significantly increased the number of macrophages (CD11b^+^F4/80^+^) and classical monocytes (GR-1^−^ CD11b^+^ LY6C^hi^) over the culture period, dependent on CSF1R signalling (**Figure 4B**). Absolute numbers of macrophages and classical monocytes significantly increased after plating, suggesting that FRCs were able to actively foster the proliferation and/or differentiation of these cells. FRCs were also able to maintain numbers of non-classical monocytes (GR-1^−^ CD11b^+^ LY6C^lo^), which otherwise underwent attrition over the culture period (**Figure 4B**).

LPS treatment, which did not alter CSF1 transcription, nonetheless had synergistic effect when administered with FRCs, increased macrophage number two-fold above the FRC-mediated increase, and this additional boost was not CSF1 dependent (**Suppl. Fig. 2B**),.

Based on these results, we explored protein-level expression of CSF1R ligands by human FRCs. In addition to expressing CSF1, human FRCs also expressed IL-34 protein (**Figure 4C, D**, confirmed by RNAseq data), which also binds CSF1R and induces the maturation of monocytes into macrophages (Foucher *et al.*, 2013).

To examine the effects of human FRCs and FRC-derived CSF1R ligands on human monocyte and macrophage phenotypes, we used a co-culture system with PBMCs from healthy donors as a monocyte source and examined numbers and phenotypes of major monocyte and macrophage subsets after 3 days. Macrophages exhibit effects on a continuum from strongly pro-inflammatory (often denoted M1; CD64^+^HLA-DR^+^) through to strongly suppressive (M2; CD206^+^CD64^−^) (Buchacher *et al.*, 2015; Tarique *et al.*, 2015). Their differentiation depends on the cues given by the local tissue micro-environment (Lenzo *et al.*, 2012). Human FRCs did not alter M1-phenotype macrophages (**Figure 4E**), but provided a strong differentiation stimulus for M2-phenotype macrophages (**Figure 4F)**. This effect of FRCs was significantly reduced in the presence of a CSF1R blocking antibody, capable of neutralizing the effects of both CSF1 and IL-34 in culture. FRCs also drove an average 8-fold increase in CD16^−^CD14^+^ classical monocytes, however this was not mediated through CSF1R signalling, since CSFR1 blockade did not prevent the increase (**Figure 4G**). As with mouse monocytes, this suggested the presence of undescribed CSF1R-independent mechanisms of FRC-support. Human FRCs did not alter numbers of CD16^+^CD14^−^ non-classical monocytes (**Figure 4H**). These data were concordant with mouse results.

We also tested the effect of LPS on this system, having observed that LPS did not alter CSF1 transcription. In the presence of LPS, FRCs still provided significant support for M2 and classical monocytes, but this occurred via CSF1R-independent mechanisms (**Figure 4F, G, Supplementary figure 2B**). Fold-change increases in M2 macrophages and non-classical monocytes with FRC co-culture represented an absolute increase in cell numbers over 72h of culture, showing that FRCs support an active increase in these cells rather than simply fostering survival (**Supplementary figure 2C, D**).

Together, these data show that mouse and human FRCs are able to promote the differentiation or expansion of M2 macrophages and classical monocytes, at least *in vitro*. FRCs expressed CSF1R ligands CSF1 and IL-34, and signaling to the CSF1R played a major role in the support observed under steady-state conditions. Additional undefined FRC-derived signals were involved in driving the FRC-mediated increase in M2 macrophages and classical monocytes that occurred in the presence of LPS.

### FRC ablation diminishes myeloid cell lineages within lymph nodes

Our data showing accumulative effects of FRCs on M2 macrophages and classical monocytes drove us to test the hypothesis that the removal of FRCs would lead to a loss of major myeloid cell types in the lymph node.

To test this, we used two mouse models of FRC depletion. *In vivo* depletion of FRCs was achieved, by crossing a CCL19-Cre strain with a Rosa26-diphtheria toxin receptor strain (CCL19-DTR), as described (Cremasco *et al.*, 2014) . In this mouse model, diphtheria toxin (DTx) treatment specifically depletes FRCs within lymph nodes. 48h after DTx treatment, lymph nodes underwent significant and prolonged involution, involving overall reduced cellularity that did not recover for the duration of the study (3 weeks) (**Figure 5A**). Accordingly, FRCs were fully ablated at day 2 post-treatment, and did not recover within the study duration (**Figure 5B**). Monocytes underwent an initial influx into the lymph node prior to their loss (**Figure 5C**), and macrophages similarly showed a delayed loss, with no effect at day 2 and a significant depletion at day 8 that did not recover by day 22 (**Figure 5D**).

**Figure 5.**
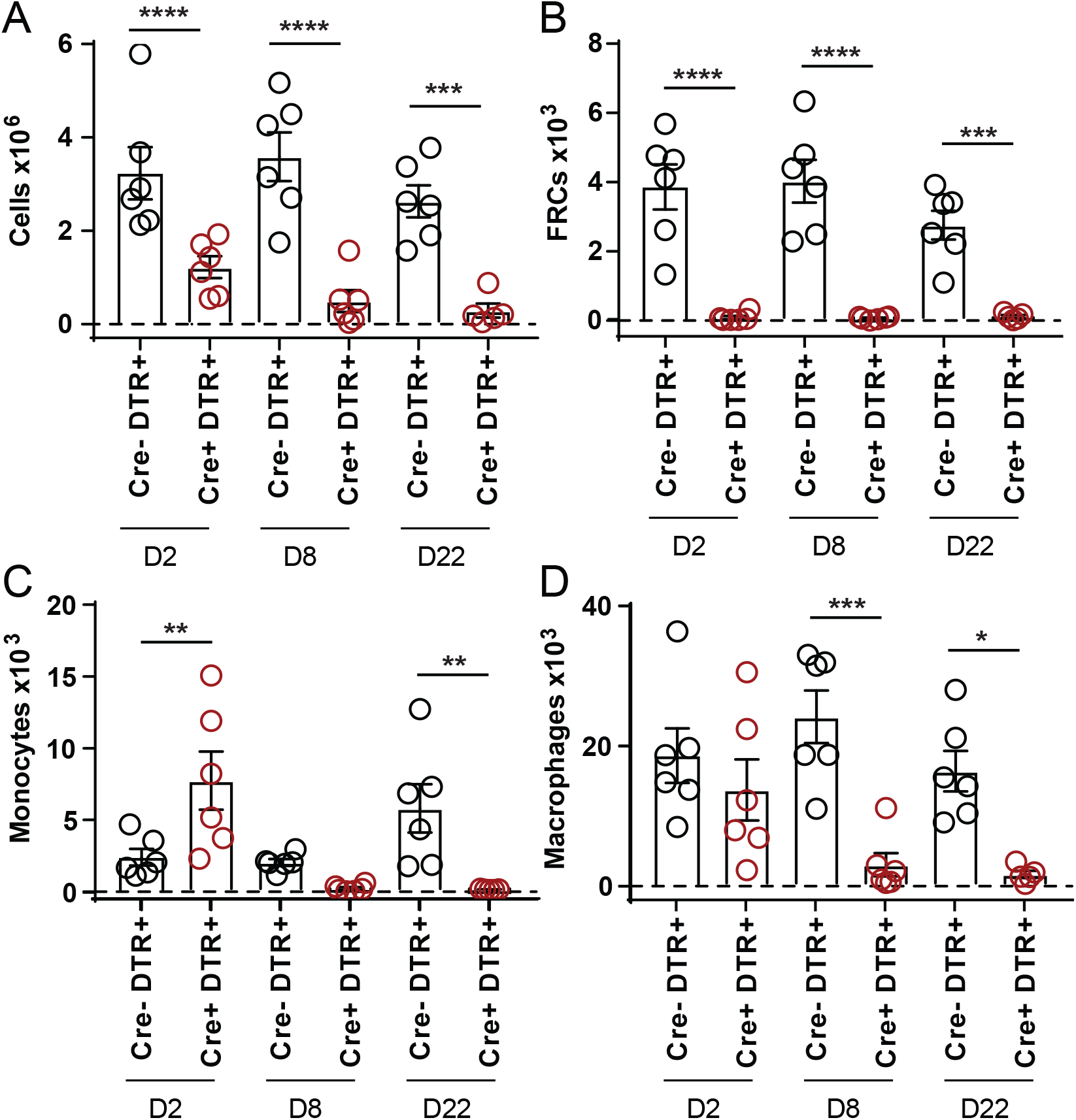
In vivo depletion of FRCs induces rapid loss of myeloid-lineage cells. CCL19-DTR mice, or DTR-expressing Cre-negative littermate controls were treated with diphtheria toxin, and brachial lymph nodes were harvested and analysed by flow cytometry at 2, 8 or 22 days after treatment ceased. **A.** Total lymph node cellularity. **B.** FRC numbers, gated as CD45^−^CD31^−^gp38^+^ cells. Monocyte numbers, gated as B220^−^NK1.1^−^CD3e^−^Ly6C^high^CD11b^+^. **D.** Macrophage numbers, gated as B220^−^NK1.1^−^CD3e^−^Ly6C^−^F480^+^. All graphs depict Mean+SEM from N=5-6 mice per group. Statistical significance was assessed using a one-way ANOVA with Sidak’s multiple comparisons test: * P<0.05, ** P<0.01, *** P<0.001, **** P<0.0001.

Monocyte and macrophage populations were therefore not sustainable in the absence of FRCs. FRCs were dispensable for a monocyte influx at Day 2, but this increase was short-lived, and did not translate to an observable recovery in macrophage numbers at timepoints studied.

In a separately developed model where FRCs express DTR under the control of Fibroblast Activation Protein (DM2 mice (Denton *et al.*, 2014)), we observed a similar effect of FRC depletion on monocyte and macrophage numbers (**Supp. Figure 3**). Lymph nodes harvested 2 days after the cessation of DTx treatment showed that FRC ablation led to a significant reduction in monocytes (classical and non-classical), and macrophages (subcapsular and medullary).

Taken together, these findings demonstrate the provision of a supportive niche by FRCs to myeloid lineage cells within secondary lymphoid organs.

## Discussion

FRCs form the structural highway on which leukocytes interact. FRCs have been shown to facilitate deletional tolerance (Fletcher *et al.*, 2010), antigen presentation (Baptista *et al.*, 2014; Dubrot *et al.*, 2014), and lymphocyte and dendritic cell homing (Bajenoff *et al.*, 2006; Link *et al.*, 2007) . FRCs promote T cell, B cell, plasma cell and ILC survival (Link *et al.*, 2007; Cremasco *et al.*, 2014; Gil-Cruz *et al.*, 2016; Huang *et al.*, 2018); they regulate T cell activation in mice and humans (Fletcher *et al.*, 2020), and are known to respond to LPS (Malhotra *et al.*, 2012; Fletcher *et al.*, 2014; Gil-Cruz *et al.*, 2016; Perez-Shibayama *et al.*, 2018). Recently, targeted deletion of type I interferon receptor (Ifnar) from FRCs revealed a role in infection-driven monocyte and neutrophil accumulation and recruitment, revealing an important biological imperative to understand FRC-innate immune cell interactions within lymph nodes and secondary lymphoid organs (Perez-Shibayama *et al.*, 2020).

While the presence of resident medullary macrophages within lymph nodes has been long reported (Bellomo *et al.*, 2018) and macrophages are confirmed in FRC-rich zones (Zhang *et al.*, 2012; Baratin *et al.*, 2017; Huang *et al.*, 2018), the cells and factors supporting their residence remain undefined. Here we sought to discover the immunoregulatory factors involved in the intimate relationship between FRCs and myeloid cells under steady-state and inflammatory conditions.

Immunofluorescence microscopy of the human tonsil and mouse lymph nodes revealed that macrophages colocalize with FRC fibers and elongate when aligned with FRC fibers. Our transcriptomic analysis of mouse and human FRCs built on previous findings and showed a strong conservation of expression of genes relevant to innate immunity, particularly the expression of CSF1, CCL2, IL-6, IL-8/CXCL8 and CXCL12, which all have well-established roles in the maturation, function, adhesion and/or chemoattraction of myeloid cells (Witmer-Pack *et al.*, 1993; Gerszten *et al.*, 1999; Pixley and Stanley, 2004; MacDonald *et al.*, 2010; Mauer *et al.*, 2014).

TLR4 (CD284), in complex with CD14 and Ly96/MD2, forms the major LPS receptor on mammalian cells, and it also binds endogenous proteins such as oxidized low-density lipoprotein and beta-defensins, as well as polysaccharides including hyaluronic acid and heparin sulfate proteoglycan (Brubaker *et al.*, 2015). TLR4 complex ligation drives multiple intracellular signalling cascades. The myeloid differentiation primary response 88 (MyD88)-dependent pathway activates the NF-κB pathway, driving transcription of pro-inflammatory cytokines, such as IL-1B and IL-6. MyD88 also activates mitogen-activated protein kinase (MAPK), which leads to activation of transcription factor AP-1, controlling gene transcription as well as mRNA stability, notably for IL-6. Some genes, such as CXCL8 which encodes IL-8, contain binding regions for both NF-κB and MAPKs, and are synergistically influenced by both (Akira and Takeda, 2004; Cronin *et al.*, 2012).

Our work showed that lymph node FRCs are poised to respond quickly to LPS-driven inflammation by upregulating innate cytokines IL-6, IL-8/CXCL8 and CCL2, through rapid TLR4 signalling, which began within 5 mins and occurred via both NF-κB and MAPK pathways. The relevance of upregulation of these cytokines *in vivo* requires further exploration, but it is reasonable to assume that production of these factors in lymph nodes following TLR signalling is very likely to stimulate resident macrophages to respond.

Single-cell RNA-seq data identified and validated eight reticular cell clusters from LPS-treated and treatment-naïve mice, correlating with previous single-cell RNA-seq analyses, and demonstrating the niche specific heterogeneity of these cell types (Rodda *et al.*, 2018; Kapoor *et al.*, 2021). From this, myeloid cell-associated chemokines, Cxcl1, Cxcl2 and Ccl2, were expressed in LPS-treated mice, suggesting an as-yet untested role for FRCs in promoting macrophage and monocyte migration. This correlates with recent findings that the recruitment of monocytes into the lymph node following immunisation with OVA-Alum is dependent on CCL2 derived from FRCs (Dasoveanu *et al.*, 2020), though these findings may be model-dependent as neither Ccl2 nor Cxcl1 expression by FRCs was upregulated in an Herpes Simplex Virus infection model, despite observed myeloid infiltration (Gregory *et al.*, 2017). Murine 3T3 fibroblasts have been shown to facilitate the migration of macrophages via cellular contractions and tunnel formation (Ford, Orbach and Rajagopalan, 2019), however the chemotactic mechanisms responsible have not been validated.

Co-culture experiments suggested that FRCs support the differentiation and survival of M2-like macrophages via signalling through CSF1R in the absence of TLR4 signalling, and through other, yet to be defined, mechanisms in the presence of LPS. It is attractive to speculate that FRCs promote macrophage polarisation towards a regenerative and repair state, and away from an inflammatory state, as FRCs dampen innate immune-driven inflammation in murine sepsis (Fletcher *et al.*, 2014; Xu *et al.*, 2019).

The importance of an intact FRC network for macrophage maintenance was demonstrated using *in vivo* depletion of FRCs in two mouse models, CCL19-DTR and FAP-DTR. Both models showed a rapid loss of resident macrophages. The lack of recovery was not unexpected; alterations in FRC networks can take weeks-to-months to resolve (Novkovic *et al.*, 2016; Gregory *et al.*, 2017). The molecular mechanisms are unknown; certainly both models induce a loss of various cell types including T cells, B cells and dendritic cells that are dependent on FRCs for survival (Cremasco *et al.*, 2014; Denton *et al.*, 2014), but there is nonetheless a clear dependence on FRCs, direct or indirect, for maintenance of monocyte and macrophage numbers within lymph nodes.

While mouse and human FRCs exhibit some clear molecular differences in regulation of T cell activation (Knoblich *et al.*, 2018), our results showed that both the effects of FRCs on macrophages and monocytes, and a CSF1R-signalling mechanism are strongly conserved between mice and humans. There was one notable difference: in mice, LPS stimulation further enhanced the capacity of FRCs to promote M2 macrophage differentiation, while in human FRC:monocyte co-cultures, it did not. The basis for this remains unclear, but differences in cell source (human PBMC vs mouse bone marrow) and markers may play a role (Murray *et al.*, 2014).

The biology of human FRCs is still largely unexplored, and it is still unclear which subset/s of FRCs are best represented through *in vitro* culture, highlighting the importance of *in vivo* follow up. Recent findings (Huang *et al.*, 2018; Rodda *et al.*, 2018; Zhang *et al.*, 2018) have shown that mouse FRC subsets include a distinct medullary population that regulates plasma cell function via production of APRIL and IL-6. Medullary macrophage populations are reportedly CSF1-independent in mice (Witmer-Pack *et al.*, 1993; Cecchini *et al.*, 1994; MacDonald *et al.*, 2010). These sinusoidal and medullary myeloid populations have recently been shown to respond to RANKL signalling from marginal reticular fibroblasts and LECs (Camara *et al.*, 2019), with LEC-derived CSF1 shown to regulate sinus-lining macrophage populations (Mondor *et al.*, 2019). T zone macrophages were relatively recently defined (Baratin *et al.*, 2017), and whether they specifically rely on CSF1 is not yet known. CSF1 and IL-34 were expressed in scRNA sequencing analysis of freshly isolated human Gremlin1^+^ fibroblasts, further supporting our in vitro microarray analysis (Kapoor *et al.*, 2021)All fibroblast subsets, including T zone subsets, expressed either CSF1 or IL-34 in mice and we demonstrated protein expression of these ligands on the whole FRC population when grown in culture. IL-34 and CSF1 both bind the CSF1R and possess similar ability to promote macrophage differentiation, though their roles diverge beyond this point, driving differential cytokine secretion by macrophages (Boulakirba *et al.*, 2018).

Using complementary mouse and human studies, our data shows that macrophages interact intimately with FRCs *in vivo* and ultimately rely on FRCs for survival. Provision of CSF1R ligands, resulting in increased survival and M2 differentiation was observed *in vitro*. FRCs are poised to swiftly respond to inflammation through TLR4 signalling, upregulating factors relevant to myeloid cell function and localization. CSF1 expression was not altered with LPS treatment, and both mouse and human FRCs switched to a CSF1R-independent mechanism of myeloid cell support in the presence of LPS. Importantly, *in vivo* depletion experiments revealed that an intact stromal network is critically important to the maintenance of macrophages and monocytes. These findings show that FRCs provide microenvironmental support to macrophages and monocytes.

### Contact for Reagent and Resource Sharing

Further information and requests for resources and reagents should be directed to and will be fulfilled by the Lead Contact, Anne Fletcher.

## Experimental Model and Subject Details

### Experimental Animals

C57BL6J mice were obtained from Monash Animal Services and housed at the Animal Research Laboratories at Monash University, Clayton. CCL19-Cre mice (Chai *et al.*, 2013) were crossed with Rosa26-iDTR mice and maintained at the Peter Doherty Institute, The University of Melbourne. DM2 mice expressing the diphtheria toxin receptor (DTR) under the regulatory elements of the murine *Fap* gene (Roberts *et al.*, 2013; Denton *et al.*, 2014) were housed at the University of Birmingham Biomedical Services Unit. For single cell RNA-Seq, BAC-transgenic C57BL/6N-Tg (Ccl19-Cre)489Biat (Ccl19-Cre) mice (Chai *et al.*, 2013) have been previously described. These mice were on the C57BL/6 genetic background, were maintained in individually ventilated cages and were used between 8 and 10 weeks of age. Experiments were performed in accordance with federal and cantonal guidelines (Tierschutzgesetz) under permission numbers SG07/19 following review and approval by the Cantonal Veterinary Office (St. Gallen, Switzerland). All mice were specific pathogen– free and cared for in accordance with institutional guidelines. All experiments received approval from relevant institutional ethics committees.

### Human tissues

Palatine tonsils were obtained from consenting donors from the National Disease Research Interchange (NDRI) resource centre or Human Biomaterials Resource Centre (HBRC), Birmingham (HTA licence 12358, 15/NW/0079), under project approval number REC/RG/HBRC/12-071. All tissues were obtained and utilised in accordance with institutional guidelines and according to the principles expressed in the Declaration of Helsinki.

## Method Details

### Lymph node digestion and FRC purification

Axillary, mesenteric and brachial lymph nodes were harvested from 5 – 10 euthanized C57BL6J mice and digested according to published protocols (Fletcher *et al.*, 2011). Human tonsils were cut into 5mm segments and enzymatically digested according to previous protocols (Fletcher *et al.*, 2011). Human or mouse cell suspensions were seeded at approximately 2900 cells/cm^2^ in αMEM complete media (Invitrogen) and cultured in 10% batch selected FBS (Sigma Aldrich) and 1% penicillin/streptomycin (Invitrogen) for 1 passage before weaning to antibiotic free conditions. Cells were harvested with 0.2% Trypsin with 5mM EDTA (Invitrogen, USA). Mouse FRCs required sorting using MACS (Miltenyi Biotec), with anti-mouse CD45 and CD31 magnetic beads, according to manufacturer’s instructions. Cells were quantified using a hemocytometer, and assessed for viability using trypan blue. Viability and purity of samples were routinely >95%.

### RNA-Seq and microarray analysis

Data from freshly isolated and cultured mouse FRCs is available at NCBI GEO accession number GSE15907 (Malhotra *et al.*, 2012) and GSE60111 (Fletcher *et al.*, 2014) and processed as described. Data from human FRCs is accessible at monash.figshare.com doi: 10.4225/03/ 5a2dae0c9b455 and was prepared and processed as described (Knoblich *et al.*, 2018). All heatmaps were generated using Morpheus matrix visualization software (Broad Institute, https://software.broadinstitute.org/morpheus).

#### Mice and LPS administration

Ccl19-Cre x R26R-EYFP mice were subcutaneously injected in one flank with 5 mg of LPS from E.coli (Sigma) together with 100 mg Ovalbumin grade VI (Sigma). Mice were sacrificed at day 3 post administration and brachial lymph nodes were removed for further lymph node stromal cell isolation and scRNA sequencing analysis.

#### Preparation of stromal cells

Brachial lymph nodes were transferred into a 24-well dish filled with RPMI 1640 medium containing 2% FCS, 20 mM Hepes (all from Lonza), 1 mg/ml Collagenase D (Sigma), 25 μg/ml DNAse I (Applichem) and Dispase (Roche). Dissociated tissues were incubated at 37°C for 30 minutes. After enzymatic digestion, cell suspensions were washed with PBS containing 0.5% FCS and 10 mM EDTA. Stromal cells were enriched by depleting CD45+ hematopoietic cells and TER119^+^ erythrocytes using MACS microbeads (Miltenyi, Germany) as described previously (Chai *et al.*, 2013). Cells were sorted for EYFP^+^CD31^−^CD45^−^ reticular cells and processed using the 10x Chromium (10X Genomics) system. Two biological replicates of LPS-treated and two biological replicates of naïve controls were used for the analysis.

#### Droplet-based single cell RNA-seq analysis

Isolated cells were sorted for EYFP^+^ reticular cells and processed on a 10x Chromium (10X Genomics) (62) to generate cDNA libraries. The processing followed the recommended protocol for the Chromium Single Cell 3’ Reagent Kit (v3 Chemistry) and the generated libraries were sequenced on an Illumina NovaSeq 6000 sequencing system at the Functional Genomic Center Zurich. Sequencing files were pre-processed using CellRanger (v3.0.2) (Zheng *et al.*, 2017) with Ensembl GRCm38.94 release as reference and damaged cells or duplicates were removed in a subsequent quality control using scater R/Bioconductor package (v1.11.2) (McCarthy *et al.*, 2017) running in R v3.6.0. In detail, cells were excluded if they had very high or low UMI counts or total number of detected genes (more than 2.5 median absolute deviations from the median across all cells) or high mitochondrial gene content (more 2.5 median absolute deviations above the median across all cells). In addition, cells expressing one of the genes Top2a, Mki67, Pclaf or Cenpf were excluded as proliferating cells resulting in a total of 10 019 cells from naïve lymph nodes and 14 427 cells from LPS treated mice.

Further downstream analysis was performed using functions from the Seurat package (v3.1.1) (Stuart *et al.*, 2019) running in R v3.6.1 including data normalization, scaling, dimensional reduction with PCA and UMAP and graph-based clustering. Clusters were characterized based on marker genes and conditions were compared based on differentially expressed genes inferred from Wilcoxon test as implemented in the FindMarkers function (Seurat v3.1.1) (Stuart *et al.*, 2019). Finally, functional differences between conditions were summarized based on a gene set enrichment analysis. For this all genes were ranked based on a signal-to-noise ratio statistic calculated on normalized expression values. Resulting ranked gene lists were used as input for GSEA-Preranked (v7.0.3) in a GenePattern notebook (Reich *et al.*, 2017) with gene sets from the mSigDB (v7.0) C2 collection. Accession number: E-MTAB-10908.

### *In vivo* FRC ablation

CCL19-Cre x Rosa26-iDTR (CCL19-DTR) mice and Cre-negative Rosa26-iDTR^+^ control mice received two injections of Diphtheria toxin (DTx) at 10ng/g of body weight, 24h apart. Skin draining LN were harvested at different days following DTx treatment and digested in RPMI containing 2% FCS, 1mg/mL collagenase D and 0.1mg/mL DNAse for 25 min at 37C. LN were further incubated for 15 min with the addition of 0.8mg/mL dispase (67). Single cell suspensions were filtered (70μm) before staining with antibodies for flow cytometry .

Diphtheria toxin receptor (DTR) FAP^+^ DM2 mice received 25ng/g diphtheria toxin (List Biological Laboratories) i.p. on days 0, 2 and 4, and were euthanised on day 6. Lymph nodes (axillary, brachial, inguinal, mesenteric) were dissected and enzymatically digested for flow cytometric analysis. Lymph node cell suspensions were labelled with cocktail containing anti-mouse conjugated antibodies. Cells were then analysed on a BD LSR Fortessa using Flowlogic software version 1.7 (Inivai Technologies).

### Luminex bead assay

Human FRCs were plated overnight at a density of 4×10^4^ cells/ 0.32cm^2^ and left to adhere to surface. Cells were then stimulated with either 1ug/ml LPS (Serotype O111:B4; Sigma Aldrich) over a 24 hour time course. Specific wells were pre-cultured for 2 hours with either 10mg/ml of TLR4 inhibitor (CLI-095, Invivogen) or 10mg/ml PI3 Kinase inhibitor (LY294002, Invivogen) as controls. Supernatants were removed at stated time points and secreted cytokine quantities were measured using Luminex Bead technology according to Bioplex protocols and cytokine Kit (Biorad).

### Phosphorylation of MAPK p38 and NFκB p65 in TLR4 stimulation

Human FRCs were plated at 4 × 10^4^ cells/ 0.32cm^2^ in complete αMEM media and left to adhere. After 4 hours, media was removed and replaced with serum free media overnight to induce basal phosphorylation levels. The following day, cells were stimulated with 1 μg/ml LPS (Serotype O111:B4; Sigma Aldrich) and cells were removed from wells at stated time points with 0.2% Trypsin with 5mM EDTA (Invitrogen), then fixed with 4% PFA and methanol, and labelled immediately according to protocols from Cell Signalling Technologies (CST) with unconjugated rabbit anti-human phosphorylated NF-κB p65 (Clone: Ser536 93H1) followed by goat anti-rabbit IgG (H+L) Alexa Fluor 488 (Thermo Fisher). For p38, the CST staining protocol was followed with mouse anti-human phosphorylated p38 MAPK (Clone: Thr180/Thr182) used with goat anti-mouse IgG (H+L) conjugated to Alexa Fluor 488 (Thermo Fisher) as a secondary. Cells were then labelled with specific stromal markers podoplanin and CD31 and were analysed using FACS Canto (BD Biosciences) with analysis undertaken using Flowlogic software version 1.7 (Inivai Technologies).

### Immunofluorescent microscopy

Murine axillary, mesenteric and brachial lymph nodes, or human tonsils, were snap-frozen in OCT (Sakura) and stored at −80C. 7μm sections were cut, air dried for 30 minutes, then fixed in chilled acetone for 20 minutes, followed by 2x PBS washes. Sections were stained with primary antibodies (**Table 1**) for 20-30 minutes in a dark humidified box at room temperature, followed by 2x 5 minute washes in PBS. Secondary antibodies were added for 20-30 minutes at room temperature. Slides were washed twice in PBS, with DAPI added to sections for 2 minutes at room temperature prior to mounting (ProLong Gold anti-fade mountant, Thermo Fisher). Slides were imaged on a Zeiss LSM 800 confocal scanning microscope.

### Cell morphology analysis

The perimeter of cells was manually drawn and the perimeter and area calculated using ImageJ software. To ensure cells were seen in cross-section, only those cells showing a DAPI^+^ nucleus were chosen for measurement. The morphology index is an inverse roundness metric calculated as perimeter^2^/4πArea, where the minimum value of 1 would be returned for a perfectly circular cell, while the further a cell deviates from circularity, the larger the values (Pinner and Sahai, 2008) .

### FRC and monocyte co-cultures

Human FRCs at passage 3 were plated overnight at 2×10^4^ cells per 0.32cm^2^ well. The following day, 4×10^5^ human PBMCs were isolated from healthy donors using Ficoll-Paque or Lymphoprep according to the manufacturer’s instructions, and added to appropriate wells. Where indicated, 10ng/ml of LPS (Cell Signalling Technology, USA), 0.1μg/ml human anti-CSF1R (Clone: 61701 – R&D systems) or anti-human isotype control (Clone: 11711 – R&D systems) were added. Cells were incubated for 72 hours at 37°C, then harvested, quantified using a Countess (Thermo Fisher), labelled with anti-human myeloid and stromal markers (**Table 1**) and analysed by flow cytometry on a LSR Fortessa (BD Biosciences).

For mouse assays, 2×10^5^ mouse FRCs were left to adhere to 6-well plates overnight. The next day 1×10^6^ mouse bone marrow cells were added to each well, with 100U/ml recombinant M-CSF (Peprotech), 10μg/ml purified anti-mouse MCSFR/CD115 (Clone: AFS98 – eBioscience) or 1μg/ml LPS (Cell Signalling Technology, USA) added as required. Some wells were pre-treated for 30 minutes with 10μg/ml TLR4 inhibitor TAK-242 (Invivogen). Cells were harvested after 4 days, quantified using a Z2 Coulter Counter (Beckman Coulter, USA) and labelled for flow cytometry then analysed on a FACS Canto (BD Biosciences) using Flowlogic software version 1.7 (Inivai Technologies).

Monocyte populations were described as classical (CD11b^+^Ly6C^hi^ in mouse, CD14^+^CD16^−^ in human) and non-classical (CD11b^+^Ly6C^low^ in mouse, CD14^−/lo^CD16^+^ in human). Macrophage populations were defined in human experiments as M1 macrophages (CD64^+^HLA-DR^+^CD206^−^) and M2 macrophages (CD64^−^CD206^+^). Mouse macrophages were defined as CD11b^+^F480^+^.

### Statistical analysis

A one-way ANOVA with Tukey’s post-test was used to compare 2 or more groups of parametric data. Mann-Whitney U test was used for non-parametric data. Normality was assessed using D’Agostino-Pearson. P< 0.05 was considered significant. To assess fold change data, a Wilcoxon rank test with a ratio paired T test was used. For pathway analysis, the top 15 upregulated genes with LPS treatment were used for KEGG analysis. Pathways shown yielded *P* values <0.05, with FDR and Benjamini-Hochberg values <0.05).

## Supporting information

STAR Methods table

## Author contributions

JD and ALF performed experiments, designed the study, analysed data and wrote the paper; TH designed the study, analysed data and wrote the paper; KK, HWC, YA, JDDC, JA, ED, performed experiments; ML, CPS and DR performed and analysed experiments; AD and SJT provided essential reagents; RB contributed essential guidance; MS designed experiments, SM and BL provided essential reagents, designed and analysed experiments.

## Acknowledgements

This work was supported by a Wellcome Trust Seed Award and a Birmingham Fellowship to AF, and by Monash University. TH was supported by a NHMRC Career Development Fellowship. The authors thank D. Fearon for usage of the DM2 mice model in this study. We gratefully acknowledge the contribution made by the University of Birmingham’s Human Biomaterials Resource Centre, which has been supported through a Birmingham Science City – Experimental Medicine Network of Excellence project.

## Declaration of interests

The authors declare no competing interests.

## Supplementary Figure legends

**Supp. Figure 1:**
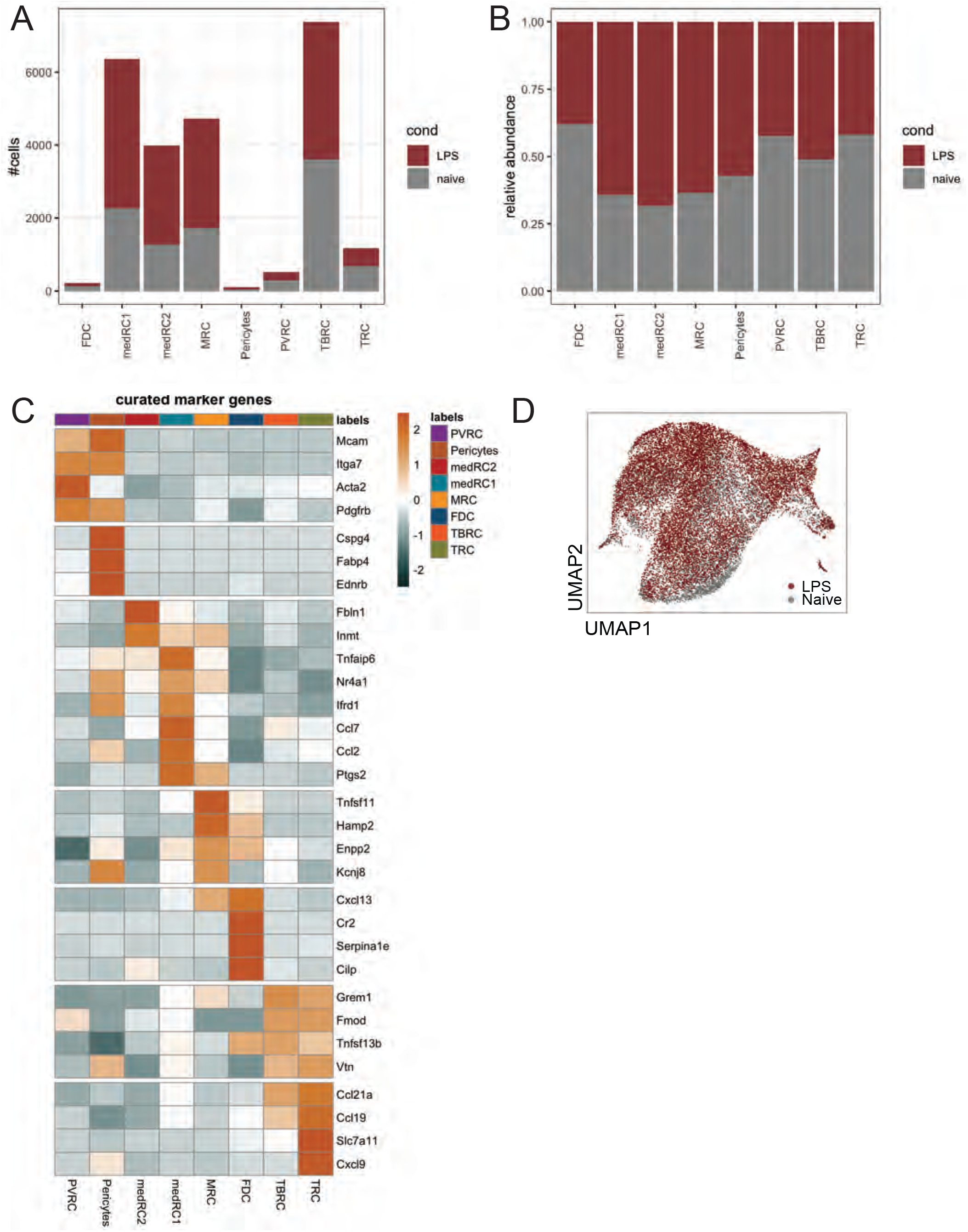
Single cell transcriptomic analysis of isolated EYFP+ reticular cells from murine lymph node stromal subsets during inflamed and resting states. EYFP^+^ reticular cells were isolated from brachial lymph nodes from Ccl19 Cre R26R-EYFP mice, which were either treatment-naïve, or immunised with OVA/LPS. scRNA-seq was performed on EYFP^+^ cells. UMAP of EYFP^+^ lymph node reticular cell subsets, categorised into 8 subsets, with or without treatment. **A, B:** Histograms of isolated EYFP+ reticular clusters showing **A.** Absolute and **B.** relative abundance of LPS treated and naïve analysed cells per reticular cell cluster. **C**. Heatmap of curated marker gene expression for identified reticular subsets. **D**. UMAP of merged data from LPS treated or naive EYFP^+^ lymph node reticular cells.

**Supp. Figure 2:**
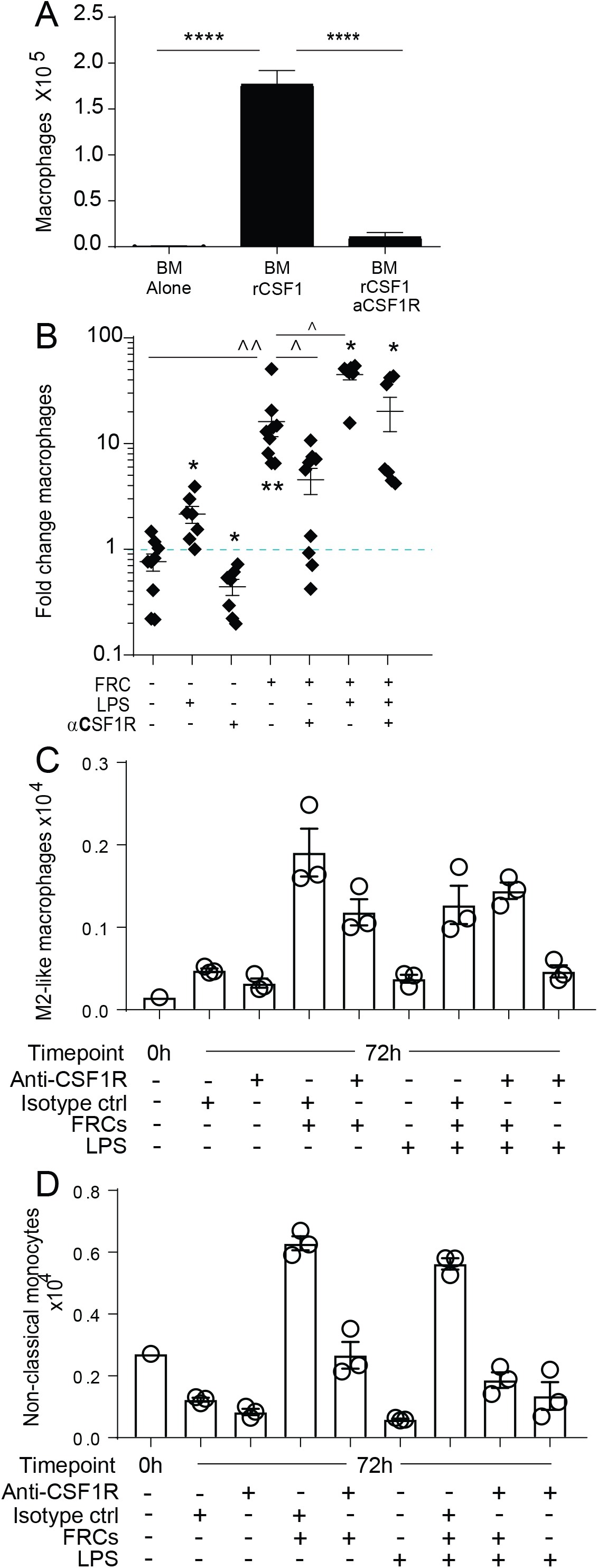
Fibroblastic reticular cells support monocyte differentiation via CSF1R signalling. 1 × 10^6^ mouse bone marrow cells, as a source of macrophage precursors, were co-cultured with 2 × 10^5^ mouse FRCs under various conditions. Cells were harvested and quantified after 3 days, quantified and assessed using flow cytometry. **A.** Macrophage numbers after 3 days of co-culture with or without recombinant CSF1 and CSF1 blocking antibody. **** P<0.0001, one-way ANOVA with Tukey’s post-test. **B.** Fold change in the number of macrophages following treatment with recombinant CSF1 and anti-CSF1R blocking antibody or LPS. The fold change shown is compared to the average of untreated controls (normalised to 1, dotted blue line). Wilcoxon rank test with a ratio paired T test was performed. Star depicts significance vs untreated group; chevron depicts significance within groups shown by horizontal lines. * or ^ P<0.05, ** or ^^ P<0.01. **C, D.** 4 × 10^5^ human peripheral blood mononuclear cells (PBMCs) were phenotyped immediately after isolation (0h) or incubated with or without LPS, CSF1R blocking antibody, isotype control antibody, or 2×10^4^ human tonsil-derived FRCs. After 72 hours of culture, cells were quantified using flow cytometry as **C.** M2-like macrophages and **D.** non-classical monocytes. Representative data from 1 FRC donor and PBMC donor is shown. For all graphs, mean + SEM shown. FRCs: fibroblastic reticular cells. CSF1: CSF1R: LPS: lipopolysaccharide. rCSF1: recombinant CSF1. αCSF1R: anti-CSF1R blocking antibody.

**Supplementary Figure 3:**
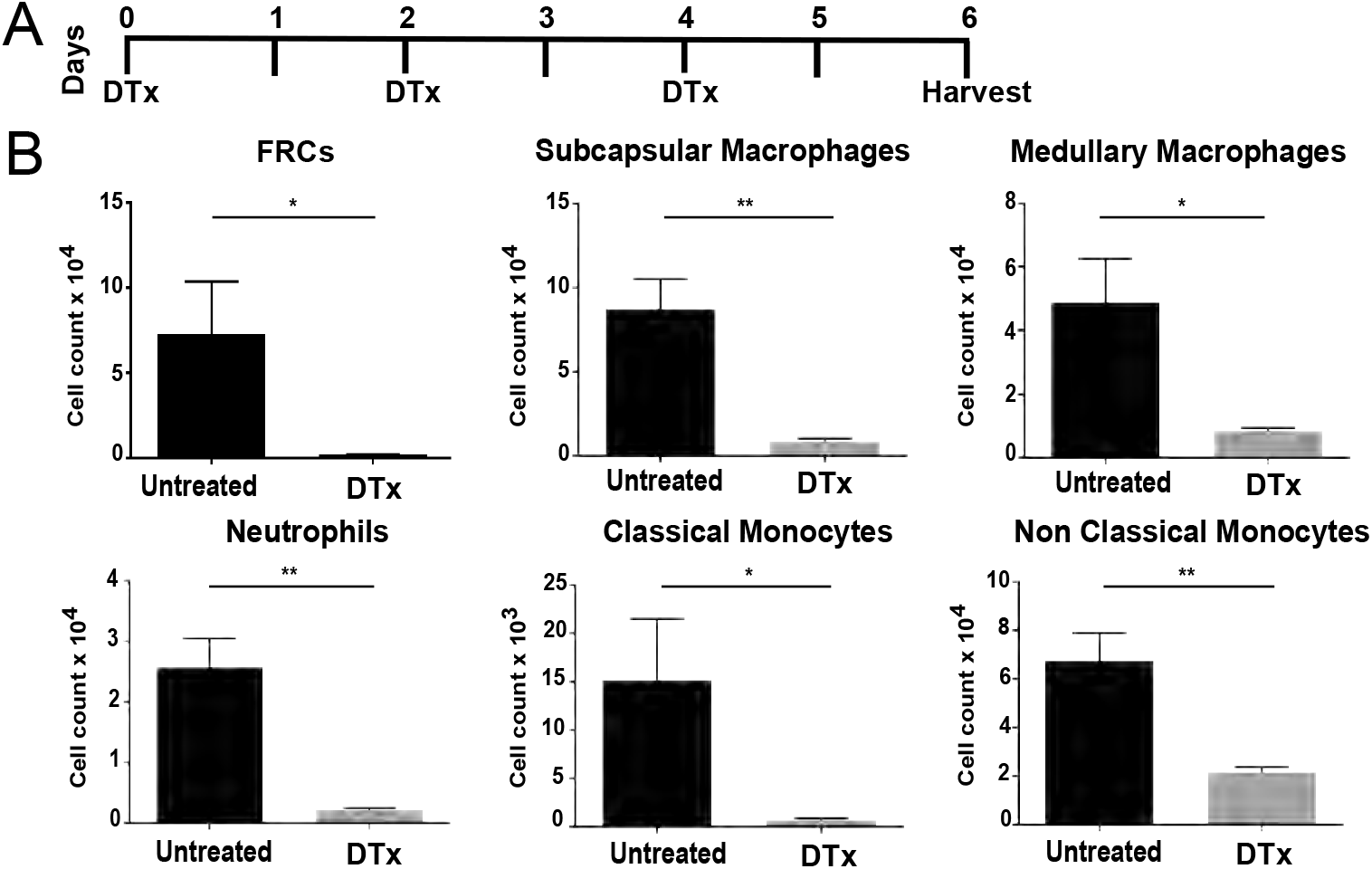
Fibroblastic reticular cell depletion leads to a reduction in innate cell populations. FAP-DTR and non-transgenic littermate control mice were each given 25ng/g diphtheria toxin (DTx) on alternate days for 6 days prior to harvest. On day 6, lymph nodes were harvested and digested according to previous protocols and analysed via flow cytometry. **A.** Timeline of DTx administration and harvest. **B.** Cell counts for lymph node macrophages (sub-capsular and medullary), neutrophils, and monocytes (classical and non-classical). Mean + SEM, n = 4 mice *P<0.05, ** P<0.01, from 1 experiment, Mann Whitney test.

## Notes

### Competing Interest Statement

The authors have declared no competing interest.

